# Genome-wide RNAi screen for regulators of UPR^mt^ in *Caenorhabditis elegans* mutants with defects in mitochondrial fusion

**DOI:** 10.1101/2020.07.31.230441

**Authors:** Simon Haeussler, Assa Yeroslaviz, Stéphane G. Rolland, Sebastian Luehr, Barbara Conradt

## Abstract

The disruption of mitochondrial dynamics has detrimental consequences for mitochondrial and cellular homeostasis and leads to the activation of the mitochondrial unfolded protein response (UPR^mt^), a quality control mechanism that adjusts cellular metabolism and restores homeostasis. To identify genes involved in the induction of UPR^mt^ in response to a block in mitochondrial fusion, we performed a genome-wide RNAi screen in *Caenorhabditis elegans* mutants lacking the gene *fzo-1*, which encodes the ortholog of mammalian Mitofusin. We find that approximately 90% of the 299 suppressors and 86 enhancers identified are conserved in humans and that one third of the conserved genes have been implicated in human disease. Furthermore, many of the 385 genes have roles in developmental processes, which suggests that mitochondrial function and the response to stress are defined during development and maintained throughout life. In addition, we find that enhancers are predominantly ‘mitochondrial’ genes and suppressors ‘non-mitochondrial’ genes, which indicates that the maintenance of mitochondrial homeostasis has evolved as a critical cellular function that when disrupted can be compensated for by a variety of cellular processes. Our analysis of ‘non-mitochondrial’ enhancers and ‘mitochondrial’ suppressors suggests that organellar contact sites, especially between ER and mitochondria, are of importance for mitochondrial homeostasis. Finally, we uncovered several genes involved in IP_3_ signaling that modulate UPR^mt^ in *fzo-1* mutants, found a potential link between pre-mRNA splicing and UPR^mt^ activation and identified the *Miga-1/2* ortholog *K01D12.6* as required for mitochondrial dynamics in *C. elegans*.

## INTRODUCTION

Mitochondria are important for cellular adenosine triphosphate (ATP) production, iron-sulfur-cluster biogenesis, lipid metabolism and apoptosis, and therefore, mitochondrial homeostasis is tightly regulated by several quality control mechanisms (Tatsuta and Langer, 2008; Kornmann, 2014). Moreover, mitochondria are required to respond to environmental challenges, which are often accompanied by alterations in energy demand (Youle and van der Bliek, 2012). Mitochondrial dynamics controls mitochondrial shape and distribution, thus playing a central role in both mitochondrial homeostasis and the adjustment to changing energy demands (Yaffe, 1999; van der Bliek *et al.*, 2013). Dynamics of mitochondrial membranes is controlled by large guanosine triphosphate-binding proteins (GTPases) of the dynamin-like family, which are conserved from yeast to humans (Hales and Fuller, 1997; Otsuga *et al.*, 1998; Smirnova *et al.*, 1998; Bleazard *et al.*, 1999; Labrousse *et al.*, 1999; Shepard and Yaffe, 1999; Chen *et al.*, 2003; Santel *et al.*, 2003; Ichishita *et al.*, 2008; Kanazawa *et al.*, 2008). In the nematode *Caenorhabditis elegans*, fusion of the outer and inner mitochondrial membrane (OMM and IMM) is facilitated by FZO-1^MFN1,2^ (Ichishita *et al.*, 2008) and EAT-3^OPA1^ (Kanazawa *et al.*, 2008), respectively. Conversely, fission of the OMM and IMM is carried out by DRP-1^DRP1^ (Labrousse *et al.*, 1999), whose ortholog in *Saccharomyces cerevisiae* (Dnm1p) has been shown to form constricting spirals around mitochondria (Ingerman *et al.*, 2005). The disruption of mitochondrial dynamics has detrimental consequences for mitochondrial and ultimately cellular homeostasis and is associated with several human diseases. Thus, mitochondrial homeostasis is controlled by several additional protective quality control mechanisms, including the mitochondrial unfolded protein response (UPR^mt^) and mitophagy (Chen and Chan, 2004; Youle and van der Bliek, 2012; van der Bliek *et al.*, 2013; Kornmann, 2014). How these quality control mechanisms are coordinated with mitochondrial dynamics is not fully understood. Recently, disruption of mitochondrial dynamics has been shown to induce UPR^mt^ (Kim and Sieburth, 2018; Zhang *et al.*, 2018; Rolland *et al.*, 2019; Haeussler *et al.*, 2020). UPR^mt^ has been studied extensively in the past decade using genome-wide RNAi screens in *C. elegans* (Haynes *et al.*, 2007; Runkel *et al.*, 2013; Bennett *et al.*, 2014; Liu *et al.*, 2014; Rolland *et al.*, 2019). Upon mitochondrial stress and the concomitant decrease in mitochondrial membrane potential, the master regulator of UPR^mt^, ‘<u>a</u>ctivating transcription factor associated with <u>s</u>tress 1’ (ATFS-1^ATF4,5^), instead of being imported into mitochondria, translocates to the nucleus, where it activates a broad transcriptional program (Haynes *et al.*, 2010; Nargund *et al.*, 2012; Rolland *et al.*, 2019). UPR^mt^ activation leads to the expression of a large set of cytoprotective genes including genes encoding chaperones (*e.g. hsp-6*^mtHSP70^ and *hsp-60*^HSDP1^, whose transcription is commonly used to monitor UPR^mt^ activation (Yoneda *et al.*, 2004)) or proteases, and has been shown to promote mitochondrial biogenesis and coordinate cellular metabolism (Nargund *et al.*, 2012; Nargund *et al.*, 2015). (All genes that are specifically up- or downregulated upon induction of UPR^mt^ are referred to as UPR^mt^ effectors.) Moreover, UPR^mt^ has been shown to act in a cell non-autonomous way and, once activated in a certain tissue, can result in a systemic response (Durieux *et al.*, 2011; Shao *et al.*, 2016; Kim and Sieburth, 2018; Zhang *et al.*, 2018; Kim and Sieburth, 2020).

In this study, we performed a genome-wide RNAi screen to identify regulators of UPR^mt^ in *fzo-1(tm1133)* mutants and identified 299 suppressors and 86 enhancers. We analyzed this dataset using bioinformatic tools, such as GO enrichment analysis, gene network analysis and analysis of transcription factor binding sites in promotors of candidate genes. Furthermore, we determined the specificities of the candidates identified with respect to their ability to modulate UPR^mt^ using secondary screens. Finally, we identified the *C. elegans* ortholog of the mammalian genes *Miga1* and *Miga2*, which have been implicated in mitochondrial fusion, and demonstrate that the loss of the *C. elegans* ortholog leads to mitochondrial fragmentation and the induction of UPR^mt^.

## METHODS

### General *C. elegans* methods and strains

*C. elegans* strains were cultured as previously described (Brenner, 1974). Bristol N2 was used as the wild-type strain. All experiments were carried out at 20°C and all strains were maintained at 20°C. The following alleles and transgenes were used: LGI: *spg-7(ad2249)* (Zubovych *et al.*, 2010); LGII: *fzo-1(tm1133)* (National BioResource Project); *eat-3(ad426)* (Kanazawa *et al.*, 2008); LGIV: *drp-1(tm1108)* (National BioResource Project); LGV: *miga-1(tm3621)* (National BioResource Project). Additionally, the following multi-copy integrated transgenes were used: *zcIs9* (P*_hsp-60_::gfp::unc-54 3’UTR*)*, zcIs13* (P*_hsp-6_::gfp::unc-54 3’UTR*) (Yoneda *et al.*, 2004); *bcIs78* (P*_myo-3_::gfp^mt^*) (Rolland *et al.*, 2013).

### RNA-mediated interference

RNAi by feeding was performed using the updated ‘Ahringer’ RNAi library (Kamath and Ahringer, 2003), which covers around ∼87% of the currently annotated *C. elegans* protein coding genes. For the primary and secondary screens with the multi-copy *zcIs13* transgene in the *fzo-1(tm1133)*, *drp-1(tm1108), eat-3(ad426)* or *spg-7(ad2249)* background, RNAi clones were cultured overnight in 100 μL of LB containing carbenicillin (100 μg/mL) in a 96 well plate format at 37°C and 200 rpm. 10 μL of each RNAi culture was used to seed one well of a 24 well RNAi plate containing 0.25% Lactose (w/v) as described previously (Rolland *et al.*, 2019). The plates were incubated at 20°C in the dark. 24 hours later, 3 L4 larvae of all strains carrying the *fzo-1(tm1133)* and *spg-7(ad2249)* allele, and 2 L4 larvae of *drp-1(tm1108)* were transferred to each well of the RNAi plates. The F1 generation was scored by eye for fluorescence intensity of the P*_hsp-6_* _mtHSP70_*gfp* reporter after 4-12 days and compared to worms of the respective genotype on the negative control *sorb-1(RNAi)*.

### Screening procedure and sequencing of RNAi-clones

For the primary screen, all RNAi clones of the library were tested once. Bacterial RNAi clones that enhanced or suppressed the P*_hsp-6_* _mtHSP70_*gfp* reporter were picked from the wells and inoculated in 100 μL of LB containing carbenicillin (100 μg/mL) in a 96 well plate format and cultured overnight at 37°C and 200 rpm. Glycerol stocks of these overnight cultures were prepared the following day by adding 100 μL of LB containing 30% glycerol and frozen at −80°C. After all RNAi clones of the library were tested, the 657 identified candidates were re-tested at least three times in duplicates for verification of the observed phenotype. The RNAi clones that reproduced the suppression or enhancement phenotype at least three out of six times were considered as verified candidates.

The 385 verified RNAi clones were sequenced. For this, colony PCRs were performed directly from the glycerol stocks using the primers *L4440F* and *L4440R*. To remove excessive primers and nucleotides, PCR products were treated with ExoSAP-IT™ (Applied Biosystems, Cat.no. 78200.200.UL) according to manufacturer’s protocol. After PCR clean-up, samples were sent for sequencing using *L4440F* primer.

*L4440 F* 5’-TGGATAACCGTATTACCGCC-3’
*L4440 R* 5’-GTTTTCCCAGTCACGACGTT-3’

According to our sequencing results, seven of the RNAi clones covered two genes. These are indicated in column B (‘Sequence’) in Table S1. These RNAi clones were assigned to the GO group of the gene, which was predominantly covered by our sequencing result and all subsequent analysis were carried out using this gene.

Subsequently, the verified and sequenced clones were rescreened in technical duplicates in three independent experiments in the secondary screens in *drp-1(tm1108)*, *eat-3(ad426)* and *spg-7(ad2249)* mutant backgrounds.

### Identification of human orthologs

Human orthologs and OMIM data (Amberger *et al.*, 2018) were extracted from wormbase.org using https://intermine.wormbase.org (Harris *et al.*, 2019). Human orthologs were then manually verified using ‘alliancegenome.org’ (The Alliance of Genome Resources, 2019), ‘orthodb.org’ (Kriventseva *et al.*, 2018), ‘ensembl.org’ (Hunt *et al.*, 2018) and ‘uniprot.org’ (Consortium, 2018).

### Prediction of mitochondrial localization and mitochondrial targeting sequences

First, https://intermine.wormbase.org (Harris *et al.*, 2019) was used to identify all candidate genes, which are related to any mitochondrial processes/pathways. To that end, we extracted all 698 genes currently associated with at least one of the 404 GO-terms containing ‘mitochond’ and checked how many of our 385 candidate genes are among them. Additionally, we used the online platform ‘MitoProt’ (https://ihg.gsf.de/ihg/mitoprot.html) (Claros and Vincens, 1996) for computational prediction of mitochondrial targeting sequences. Proteins for which the prediction of a mitochondrial targeting sequence was ≥0.5 were considered to be mitochondrial.

### Gene ontology enrichment analysis using DAVID

In search of enriched gene ontology terms, we used the DAVID tool (version 6.8 (Huang *et al.*, 2008, 2009)) and ran the list of candidates against all genes of the *C. elegans* genome as a background list. Using an EASE score from the modified fisher-exact test, the clustering algorithm groups genes based on their association in GO categories and assigns a significance value to the group (Huang *et al.*, 2007). The clustered groups were then plotted using modified functions from the GO plot package (R version 1.0.2 (Walter *et al.*, 2015)).

### Transcription factor enrichment analysis

We searched for enriched transcription factors using the tool g:Profiler (a tool for functional enrichment analysis using over-representation (Raudvere *et al.*, 2019)). The two input lists (suppressors and enhancers of *fzo-1(tm1133)*-induced UPR^mt^) with WBGene-IDs of the identified candidate genes were used to search in the Transfac database (annotations: TRANSFAC Release 2019.1 classes: v2 (Knüppel *et al.*, 1994; Matys *et al.*, 2006)).

### Construction of gene networks of FZO-1 and MFN1/2, and the UPR^mt^

The *C. elegans* interactomes were compiled for FZO-1 or all 16 genes that are currently associated with the GO-term ‘mitochondrial unfolded protein response’ (GO:0034514) from scientific literature (Durinck *et al.*, 2009; Simonis *et al.*, 2009) and databases such as mentha (Calderone *et al.*, 2013), BioGRID3.5 (Oughtred *et al.*, 2018), IntAct (Orchard *et al.*, 2014) and STRING (Szklarczyk *et al.*, 2018) (STRING was only used to build the FZOome). The human orthologs of those genes were identified and were searched as well. Whenever possible, the interaction partners were converted back to *C. elegans* genes using biomaRt (Durinck *et al.*, 2009) and available scientific literature (Shaye and Greenwald, 2011; Kim *et al.*, 2018). The complete list of interactions was uploaded to cytoscape (v.3.7.2 (Shannon *et al.*, 2003)) and a network was calculated, highlighting both enhancers and suppressors from the screening results.

### Image acquisition, processing and analysis

For each mutant (Figure S1), 10-20 animals were immobilized with M9 buffer containing 150 mM sodium azide on 2% agarose pads and imaged using a Leica GFP dissecting microscope (M205 FA) and Leica Application Suite software (3.2.0.9652).

Mitochondrial morphology was assessed in a strain carrying *bcIs78* (*P_myo-3_::gfp^mt^*) using a Zeiss Axioskop 2 with a 63x objective and MetaMorph software (Molecular Devices).

## RESULTS & DISCUSSION

### Genome-wide RNAi screen for suppressors and enhancers of *fzo-1(tm1133)*-induced UPR^mt^ identifies highly conserved set of genes with relevance to human health

The disruption of mitochondrial dynamics in *C. elegans* induces the mitochondrial unfolded protein response (UPR^mt^) (Kim and Sieburth, 2018; Zhang *et al.*, 2018; Rolland *et al.*, 2019; Haeussler *et al.*, 2020). To identify genes affecting mitochondrial homeostasis in animals with defects in mitochondrial dynamics, we used a loss-of-function mutation of *fzo-1^MFN1,2^, tm1133*, (National BioResource Project) to induce the UPR^mt^ reporter P*_hsp-6_* _mtHSP70_*gfp (zcIs13)* and screened the *C. elegans* genome for modifiers. To that end, we used RNA-mediated interference (RNAi) and targeted ∼87% of the currently annotated protein coding genes (Kamath and Ahringer, 2003) (Figure 1A). The moderate induction of the P*_hsp-6_* _mtHSP70_*gfp* reporter in the *fzo-1(tm1133)* background allowed the identification of both suppressors and enhancers of the response. Using a protocol in which the F1 generation is scored for P*_hsp-6_* _mtHSP70_*gfp* expression levels in the fourth larval stage of development (L4), we initially identified 657 candidate genes of which 385 reproduced. Of the 385 candidates identified, 299 act as suppressors and 86 as enhancers (Figure 1B and Table S1). In order to assess whether the 86 identified enhancers are specific to the *fzo-1(tm1133)* background or if their depletion induces UPR^mt^ also in the absence of mitochondrial stress, we knocked them down in a wild-type background and tested for induction of the P*_hsp-6_* _mtHSP70_*gfp* reporter. All except three genes (*copd-1*^ARCN1^, *F25H9.6*^PPCDC^, *F32A7.4*^METTL17^) induce P*_hsp-6_* _mtHSP70_*gfp* expression when knocked-down in wild-type animals, suggesting that the induction of UPR^mt^ by depletion of these candidates is independent of the loss of *fzo-1*. (Candidates that encode mitochondrial proteins and that induce UPR^mt^ in a wild-type background upon knock-down were included in a recent publication, which reported the systematic identification of mitochondrial inducers of UPR^mt^ (Rolland *et al.*, 2019)).

**Figure 1:**
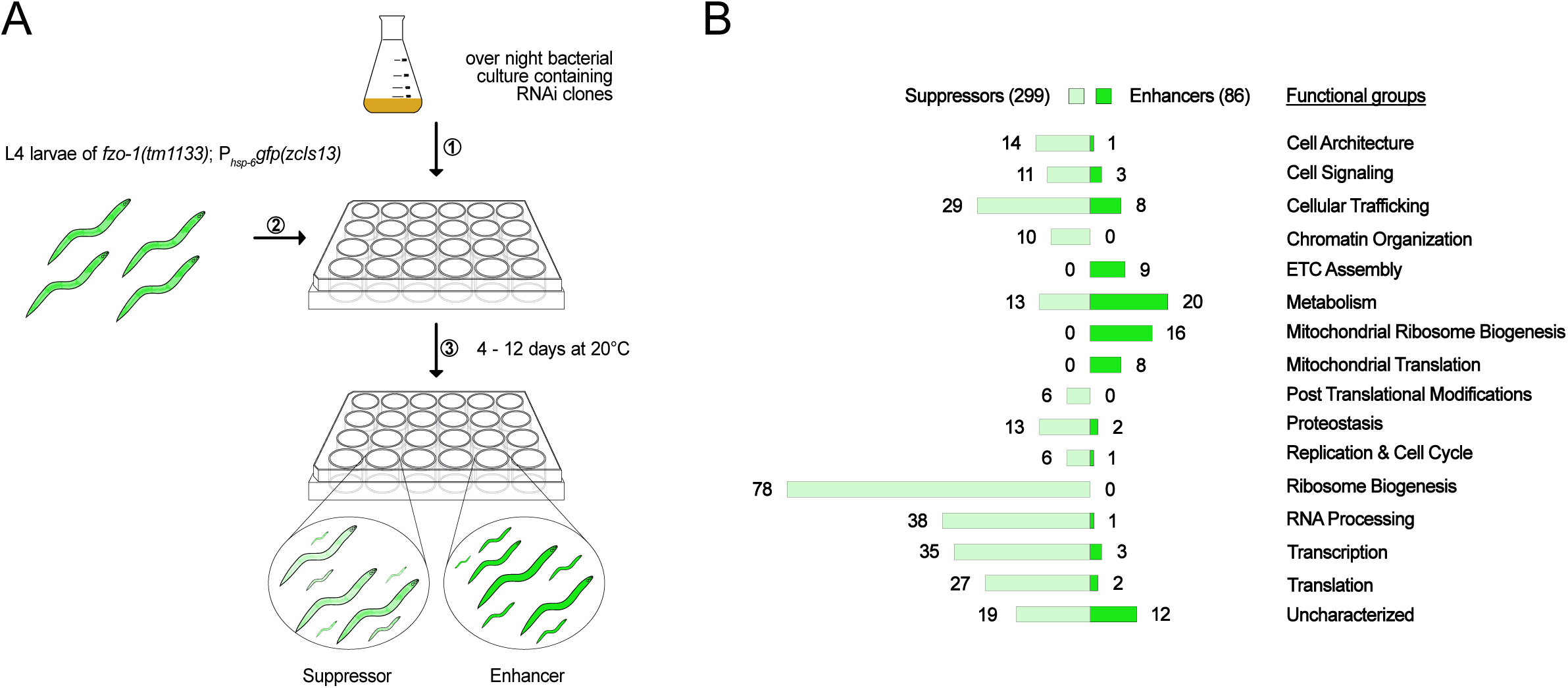
Overview of genome-wide RNAi screen for suppressors and enhancers of *fzo-1(tm1133)*-induced UPR^mt^. **(A)** Schematic overview of the RNAi screening procedure using the RNAi feeding library (Kamath and Ahringer, 2003) in *fzo-1(tm1133)* mutants that express the UPR^mt^ reporter P*_hsp-6_* _mtHSP70_*gfp (zcIs13)*. The moderate induction of the reporter in the *fzo-1(tm1133)* background allowed screening for both suppressors and enhancers of the response. **(B)** The screen resulted in identification of 299 suppressors and 86 enhancers of *fzo-1(tm1133)*-induced UPR^mt^, which were sorted into categories that we defined according to their function. ETC: electron transport chain.

Among the 299 suppressors, only 25 (8%) have previously been found to suppress UPR^mt^ induced by other means upon knock-down (Haynes *et al.*, 2007; Runkel *et al.*, 2013; Liu *et al.*, 2014). Similarly, among the 86 enhancers, only 15 (17%) have previously been shown to induce UPR^mt^ upon knock-down (indicated ‘previously identified’ in the ‘overview’ sheet of Table S1). This may be due to different genetic backgrounds and to differences in RNAi-protocols. Moreover, false negatives in RNAi screens have been estimated to vary between 10% and 30%, even if the same protocol is used by the same laboratory (Simmer *et al.*, 2003).

Using ‘alliancegenome.org’ (The Alliance of Genome Resources, 2019), ‘orthodb.org’ (Kriventseva *et al.*, 2018), ‘ensembl.org’ (Hunt *et al.*, 2018), ‘uniprot.org’ (Consortium, 2018) and ‘wormbase.org’ (Harris *et al.*, 2019) databases, we found that approximately 90% of the suppressors and enhancers (348/385) have at least one ortholog in humans (indicated ‘Human ortholog’ in the ‘overview’ sheet of Table S1). For comparison, the overall conservation of genes from *C. elegans* to humans is only about 41% (Shaye and Greenwald, 2011; Kim *et al.*, 2018). Moreover, we found that the orthologs of 36% of the conserved candidates (126/348) have previously been associated with human disease and are listed in the ‘Online Mendelian Inheritance in Man’ database (Amberger *et al.*, 2018) (indicated ‘OMIM’ in the ‘overview’ and ‘OMIM’ sheet of Table S1). In summary, we identified a set of predominantly conserved genes, many of them relevant to human health, which when knocked-down affect mitochondrial homeostasis in mutants with defects in mitochondrial fusion.

### Genes with functions in development, receptor-mediated endocytosis and metabolism modulate UPR^mt^ signaling

In order to obtain an overview of the type of processes that affect *fzo-1(tm1133)-*induced UPR^mt^, we analyzed the gene ontology (GO) terms of all 385 candidates, sorted them into ‘functional groups’ (Figure 1B) and performed a clustered gene enrichment analysis using DAVID (Huang *et al.*, 2008, 2009) (Table S2 and Figure 2). (Of note, 31 suppressors and enhancers could not be assigned to functional groups since these genes are uncharacterized in *C. elegans* and/or lack orthologs in humans. For this reason, they were assigned to the functional group ‘uncharacterized’ (Figure 1B)).

**Figure 2:**
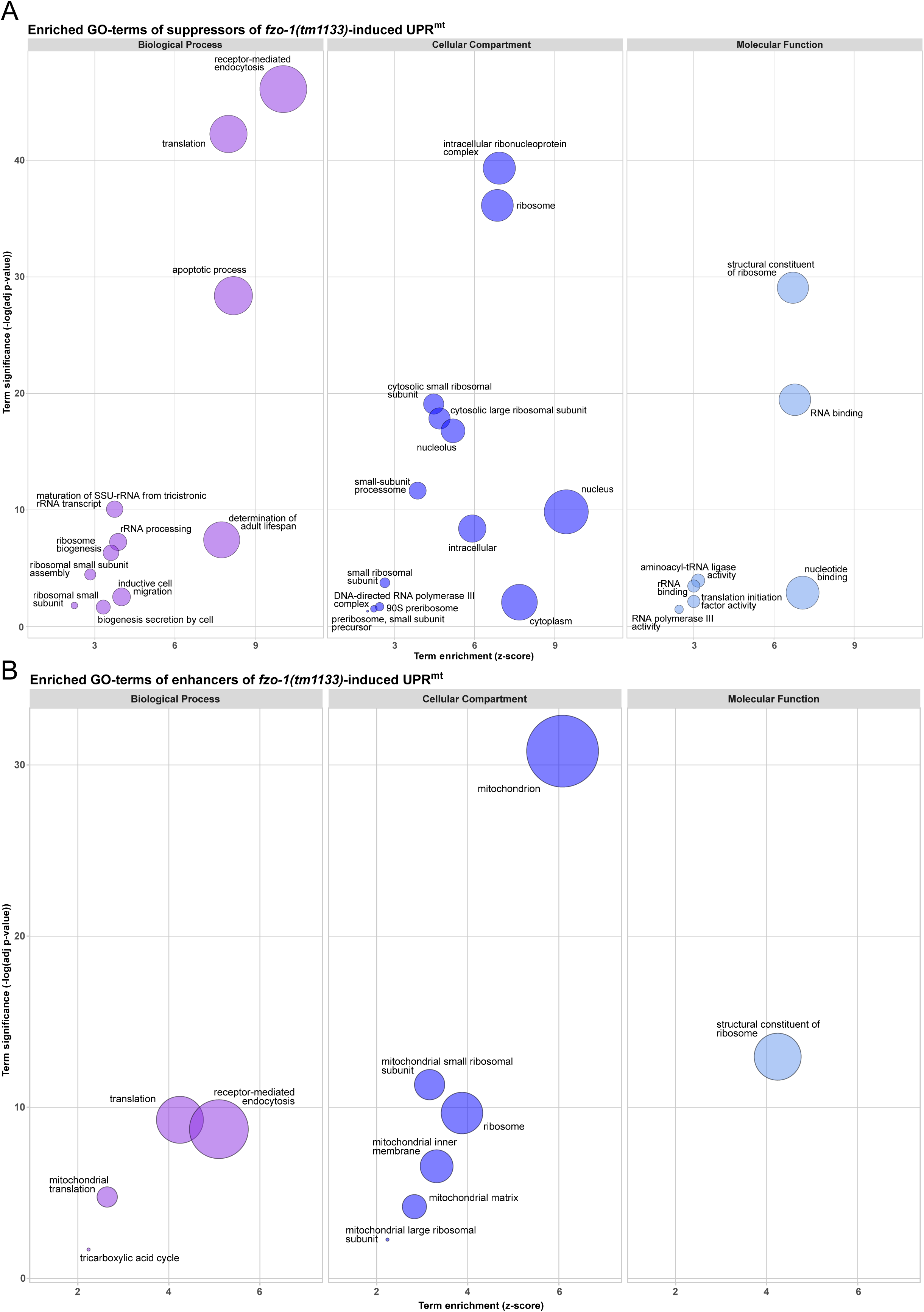
Gene ontology enrichment analysis of suppressors and enhancers of *fzo-1(tm1133)*-induced UPR^mt^ using DAVID. **(A)** Results of the clustered gene ontology enrichment analysis of suppressors of *fzo-1(tm1133)*-induced UPR^mt^ using DAVID (Huang *et al.*, 2008, 2009). **(B)** Results of the clustered gene ontology enrichment analysis of enhancers of *fzo-1(tm1133)*-induced UPR^mt^ using DAVID. **(A) & (B)** Statistically significant (*P* >0.05) enriched GO-terms, except the nematode specific GO-terms, of *fzo-1(tm1133)*-induced UPR^mt^ are depicted. Circle size correlates with the number of genes associated with a specific GO-term.

In the clustered gene enrichment analysis, we found that the majority of both suppressors and enhancers are associated with at least one of the following GO-terms: ‘nematode larval development’, ‘embryo development ending in birth or egg hatching’ or ‘reproduction’ (Table S2). It has been shown that reducing the functions of some genes encoding components of the ETC (e.g. *cox-5B(RNAi)*) in specific tissues and at specific times during development can lead to both systemic activation of UPR^mt^ and longevity (Dillin *et al.*, 2002; Rea *et al.*, 2007; Durieux *et al.*, 2011). This indicates that the activity levels of mitochondria in an individual animal are ‘set’ at a specific developmental stage and, once set, are maintained throughout development and adult life. Our results demonstrate that disrupting development compromises this process, thereby affecting an animal’s ability to cope with mitochondrial stress and to respond to UPR^mt^ activation, which will indirectly influence processes such as its lifespan. In support of this notion, we found that approximately 20% of the suppressors carry the GO-term ‘determination of adult lifespan’.

Among the suppressors, the GO-term ‘receptor-mediated endocytosis’ is enriched (Figure 2A and Table S2). It contains many genes with roles in vesicular trafficking and vesicle budding. Genes required for vesicular trafficking have been shown to affect mitochondrial morphology and homeostasis when inactivated, and it has been proposed that this is the result of altered contact sites between organelles and altered lipid transfer into mitochondria (Altmann and Westermann, 2005). Furthermore, we recently demonstrated that approximately half of the candidates in this GO-category are negative regulators of autophagy. Upon knock-down, these genes suppress *fzo-1(tm1133)*-induced UPR^mt^ most probably by inducing autophagy thereby causing changes in lipid metabolism (Haeussler *et al.*, 2020). Moreover, many cellular signaling pathways originate at the plasma membrane and, thus, are dependent on endocytosis (Sorkin and von Zastrow, 2009; Di Fiore and von Zastrow, 2014). Therefore, we speculate that depletion of genes associated with the GO-term ‘receptor-mediated endocytosis’ may either cause changes in lipid metabolism thereby suppressing UPR^mt^ or disrupt cell non-autonomous UPR^mt^ signaling.

The functional group ‘ribosome biogenesis’ contains 78 (26%) of the suppressors (Figure 1B) and includes both small- and large ribosomal subunits, as well as proteins with roles in the maturation or transport of ribosomal subunits and rRNAs. Accordingly, in all three GO-domains (Biological Process, Cellular Compartment, Molecular Function), we found that several GO-terms related to the ribosome were significantly enriched (Figure 2A and Table S2). (The GO-term ‘apoptotic process’ also contains many ribosomal subunits resulting in its enrichment in our analysis.) Moreover, we assigned a substantial part of the suppressors to the groups ‘RNA processing’ (38), ‘transcription’ (35) and ‘translation’ (27) (Figure 1B). Hence, we found five GO-terms related to translation-, two to transcription- and one to RNA-related processes to be enriched in a statistically significant manner in the GO enrichment analysis (Figure 2A and Table S2). These results raise the question whether knock-down of candidates involved in cytosolic translation specifically suppresses UPR^mt^ or simply reduces the expression of the P*_hsp-6_* _mtHSP70_*gfp* reporter. We recently showed that knock-down of the cytosolic tRNA synthetase *hars-1*^HARS1^, which we found to suppress P*_hsp-6_* _mtHSP70_*gfp* expression in *fzo-1(tm1133)* and which presumably compromises cytosolic translation, results in reduced expression of a control reporter, P*_ges-1_* _GES2_*gfp* (Haeussler *et al.*, 2020). Therefore, we cannot exclude the possibility that the knock-down of candidates related to the functional groups of transcription, RNA processing, ribosome biogenesis and translation may, at least to some extent, interfere with reporter expression *per se*.

Among the enhancers, we assigned most candidates to the functional groups ‘metabolism’ and ‘mitochondrial ribosome biogenesis’ as well as ‘cellular trafficking’, ‘mitochondrial translation’ and ‘ETC assembly’ (Figure 1B). Accordingly, GO analysis of the enhancers shows that the cellular compartments ‘mitochondrion’, ‘mitochondrial small ribosomal subunit’, ‘mitochondrial large ribosomal subunit’, ‘mitochondrial inner membrane’, ‘mitochondrial matrix’ and ‘ribosome’ are enriched (Figure 2B and Table S2). In addition, the biological processes ‘translation’ (which also includes ‘mitochondrial translation’), ‘tricarboxylic acid cycle’ and ‘receptor-mediated endocytosis’ are enriched as is the molecular function ‘structural constituent of ribosome’ (Figure 2B and Table S2). Among the enhancers carrying the GO-term ‘receptor-mediated endocytosis’, we identified many subunits of the mitochondrial ribosome and genes required for mitochondrial translation, which are most likely mis-annotated and therefore led to enrichment of this GO-term. In summary, we show that disrupting mitochondrial translation and metabolism enhances UPR^mt^ in *fzo-1(tm1133)*. Disruption of these processes has previously been shown to induce UPR^mt^ in wild type (Durieux *et al.*, 2011; Houtkooper *et al.*, 2013). Therefore, reducing mitochondrial function induces UPR^mt^ independently of the genetic background.

In summary, the GO enrichment analysis revealed that depletion of the majority of candidates in our dataset may modulate UPR^mt^ due to their role in development. Furthermore, we propose that suppressors with roles in endocytosis modulate UPR^mt^ signaling indirectly and speculate that cellular signaling and/or alterations in organellar contact sites may influence mitochondrial metabolism and hence, UPR^mt^ signaling. Finally, we find disruption of mitochondrial metabolism and translation to robustly enhance UPR^mt^ signaling in *fzo-1(tm1133)*.

### Mitochondrial fitness balances cellular homeostasis

Next, we determined which fraction of the identified enhancers and suppressors encode proteins that have a mitochondrial function or localize to mitochondria. We extracted all 698 genes that are associated with at least one of the 404 GO-terms containing ‘mitochond’ using the ‘WormMine’ database (https://intermine.wormbase.org) (Harris *et al.*, 2019), and then determined how many of our candidate genes are associated with any of these GO-terms. Using this approach, we identified 11 suppressors and 59 enhancers that encode proteins that localize to mitochondria or play a role in mitochondrial metabolism and dynamics, respectively (indicated ‘GO mitochond’ in ‘Overview’ and ‘Mitochondrial’ sheet of Table S1). Next, we used the online platform ‘MitoProt’ (https://ihg.gsf.de/ihg/mitoprot.html) (Claros and Vincens, 1996) for computational prediction of mitochondrial targeting sequences and identified an additional 5 suppressors and 14 enhancers that are predicted to localize to mitochondria (cut-off value ≥ 0.5) (indicated ‘MitoProt prediction’ in ‘Mitochondrial’ sheet of Table S1). Third, by literature searches, we found that the orthologs of 3 enhancers localize to mitochondria (Shafqat *et al.*, 2003; Spaan *et al.*, 2005; Cambier *et al.*, 2012). In summary, 76 out of 86 (88%) enhancers and 16 out of 299 (5%) suppressors encode proteins that have a mitochondrial function. This suggests that only few processes exist outside of mitochondria that can perturb mitochondrial homeostasis when disrupted. Conversely, many processes and mechanisms exist outside of mitochondria that can compensate for mitochondrial dysfunction, thereby ensuring mitochondrial and consequently cellular homeostasis.

Among the 10 ‘non-mitochondrial’ enhancers of UPR^mt^ are three genes (*F29B9.8*, *Y61A9LA.11*, *C25H3.10*) with yet unknown function, which lack orthologs in other systems. ORC-1^ORC1^ is a component of the origin recognition complex and plays a role in DNA replication (Gavin *et al.*, 1995; Ohta *et al.*, 2003; Tatsumi *et al.*, 2003). The disruption of DNA replication or cell cycle progression has previously not been reported to lead to UPR^mt^ induction. We speculate that disruption of DNA replication leads to developmental defects and therefore induces UPR^mt^. *F25H9.6*^PPCDC^ is the *C. elegans* ortholog of phosphopantothenoylcysteine decarboxylase, an enzyme required for the biosynthesis of coenzyme A (CoA) (Daugherty *et al.*, 2002). Thus, knock-down of *F25H9.6*^PPCDC^ may interfere with critical biosynthetic and metabolic pathways (including the TCA cycle) and therefore enhance UPR^mt^. NHR-209^HNF4A,G^ is orthologous to Hepatocyte Nuclear Factor 4α (HNF4A) and belongs to the family of nuclear hormone receptors, a class of cofactor and ligand-inducible transcription factors (TFs) that regulate various cellular processes, including metabolism, development and homeostasis (Aranda and Pascual, 2001; Bolotin *et al.*, 2010). Interestingly, long-chain fatty acids are ligands of HNF4A and, depending on their chain length and degree of saturation, activate or repress the transcriptional activity of HNF4A (Hertz *et al.*, 1998; Dhe-Paganon *et al.*, 2002; Wisely *et al.*, 2002; Duda *et al.*, 2004). Furthermore, HNF4α activity has been shown to be required for ß-oxidation of fatty acids both in mice and *Drosophila melanogaster* (Palanker *et al.*, 2009; Chen *et al.*, 2020). Thus, NHR-209^HNF4A,G^ may have a similar role in *C. elegans* and act as a metabolic sensor, which when deactivated enhances UPR^mt^ in *fzo-1(tm1133)*. Moreover, we identified *cpna-3*^CPNE5,8,9^, an ortholog of mammalian copine family members, a class of calcium dependent phospholipid binding proteins with roles in intracellular signaling and membrane trafficking (Creutz *et al.*, 1998; Tomsig *et al.*, 2003; Tomsig *et al.*, 2004; Ramsey *et al.*, 2008). Previously, another gene of the copine family, *gem-4*^CPNE8^, was shown to be upregulated upon UPR^mt^ activation (Nargund *et al.*, 2012). Therefore, we speculate that signaling via copine family members may be important for UPR^mt^ regulation. Another non-mitochondrial enhancer, *copd-1*^ARCN1^, encodes a protein orthologous to the delta subunit of coatomer in *S. cerevisiae* and humans (RET2 and ARCN1, respectively), which is involved in the formation of coat protein complex I (COPI) vesicles. COPI vesicles play a central role in the secretory pathway and are required for the retrieval of lipids and proteins from the Golgi apparatus and their subsequent retrograde transport to the ER (Lee *et al.*, 2004; Beck *et al.*, 2009). Furthermore, the trafficking to their final destination of most non-mitochondrial and non-peroxisomal transmembrane proteins, as well as proteins required for the release of neurotransmitters, such as SNARE proteins, is dependent on COPI-mediated transport (Beck *et al.*, 2009). Thus, disruption of the secretory pathway affects many intra- and intercellular signaling pathways, including the Ras and TOR signaling pathways, as well as signaling via G-protein-coupled receptors (GPCRs) and receptor tyrosine kinases (Farhan and Rabouille, 2011). Moreover, disruption of the retrograde transport system has been shown to lead to erroneous secretion of ER resident proteins (e.g. ER chaperones) and, consequently, to the activation of UPR in the ER (UPR^ER^) (Aguilera-Romero *et al.*, 2008; Izumi *et al.*, 2016). Therefore, we speculate that the enhancement of UPR^mt^ induction in *fzo-1(tm1133)* animals upon *copd-1(RNAi)* may be due to alterations in one of the above-mentioned signaling pathways. This notion is supported by the finding that phospholipase C (PLC-1^PLCE1^), a GPCR associated enzyme, is among the non-mitochondrial enhancers, as well as *srh-40* (<u>s</u>erpentine <u>r</u>eceptor class <u>H</u>), which is predicted to encode a GPCR. Taken together, we identified many genes among the ‘non-mitochondrial’ enhancers, which regulate intra- and intercellular signaling cascades, and we speculate that these may play a role in signaling of UPR^mt^ both in a cell autonomous and cell non-autonomous fashion. In addition, we identified ‘non-mitochondrial’ enhancers that directly regulate metabolic homeostasis and, thus, enhance UPR^mt^ in *fzo-1(tm1133)* mutants.

Among the 16 identified ‘mitochondrial suppressors’ of UPR^mt^ are candidates, such as TFG-1^TFG^ and GBF-1^GBF1^, which encode proteins that have been shown to associate with mitochondria but also other organelles. GBF-1^GBF1^ is a guanine nucleotide exchange factor (GEF) for the small GTPase ARF-1.2^ARF1^, which in yeast recruits ARF-1.2^ARF1,3^ to ER-mitochondria contact sites (Ackema *et al.*, 2014). Depletion of GBF-1^GBF1^ leads to altered ARF-1.2^ARF1,3^ localization and changes in mitochondrial morphology both in yeast and *C*. *elegans,* and this appears to be independent of their roles in endosomal transport (Ackema *et al.*, 2014). Ackema and colleagues observed an increase in mitochondrial connectivity upon GBF-1^GBF1^ depletion, similar to that observed upon knock-down of *miro-1*^MIRO1^ and *vdac-1*^VDAC^, both of which encode proteins that also localize to ER-mitochondria contact sites. However, the alterations in mitochondrial morphology of FZO-1^MFN1,2^ depleted animals were shown to be epistatic to the changes in mitochondrial morphology observed upon *gbf-1(RNAi)* and *arf-1.2(RNAi)*. Therefore, the suppression of UPR^mt^ observed in *fzo-1(tm1133)* animals upon *gbf-1(RNAi)* may not be due to a rescue of the mitochondrial morphology defect but rather be the consequence of changes in ER-mitochondria contact sites. This highlights the importance of organellar contact sites for the maintenance of mitochondrial and consequently cellular homeostasis. Furthermore, we identified TFG-1^TFG^, a component of the secretory pathway via COPII vesicles (Witte *et al.*, 2011), as a suppressor of *fzo-1(tm1133)*-induced UPR^mt^. COPII vesicles transport newly synthesized proteins and lipids from specialized ER zones, so called ER exit sites (ERES), to the Golgi apparatus (Budnik and Stephens, 2009; Kurokawa and Nakano, 2018). Similar to what we propose for *copd-1(RNAi)* (see above), we speculate that disruption of the secretory pathway may lead to alterations in cellular signaling, ER-mitochondria contact sites and, depending on the context, either to suppression or enhancement of UPR^mt^.

Taken together, we demonstrate that the perturbation of primarily mitochondrial processes leads to the enhancement of UPR^mt^. However, the identification of non-mitochondrial enhancers demonstrates that disruption of processes taking place outside of mitochondria can also compromise mitochondrial function and activate or enhance UPR^mt^. Alterations in cellular signaling pathways and/or organellar contact sites may play a role in this respect. Moreover, we find that the majority of suppressors of *fzo-1(tm1133)*-induced UPR^mt^ are non-mitochondrial, suggesting that many cellular pathways outside of mitochondria exist that can compensate for mitochondrial stress and, hence, ensure mitochondrial homeostasis. In line with this notion, we identified a few ‘mitochondrial suppressors’, most of which are involved in the maintenance of contacts to other organelles, especially the ER.

### Defects in mitochondrial fusion and fission are suppressed and enhanced by the same pathways

In order to define the specificity of the 299 suppressors and 86 enhancers, we carried out secondary screens. To identify general modifiers of UPR^mt^, we rescreened the candidates in the background of *spg-7(ad2249),* which induces UPR^mt^ (Figure S1). *spg-7*^AFG3L2^ encodes a mitochondrial matrix AAA-protease, which induces UPR^mt^ when depleted and which is commonly used as a positive control for UPR^mt^ activation (Yoneda *et al.*, 2004; Haynes *et al.*, 2007; Haynes *et al.*, 2010). To identify genes in our dataset that specifically modify UPR^mt^ induced by defects in mitochondrial membrane fusion, we rescreened all candidates in the *eat-3(ad426)* background, in which IMM fusion is blocked. Finally, to identify genes that may modulate UPR^mt^ induced by defects in mitochondrial dynamics, we rescreened all candidates in the *drp-1(tm1108)* background, in which mitochondrial fission is blocked. In the *drp-1(tm1108)* background, of the 385 candidates, 290 suppress and 53 enhance. In the *eat-3(ad426)* background, 242 suppress and, 48 enhance. Finally, in the *spg-7(ad2249)* background, 181 suppress and 47 enhance (Table S1). (Of note, there is an inverse correlation between the level of P*_hsp-6_* _mtHSP70_*gfp* expression in the different mutant backgrounds and the number of candidates that reproduce. Hence, the level of reporter expression may correlate with the number of false negatives in a given dataset of the secondary screens, for both suppressors and enhancers.) Since more suppressors reproduced in *drp-1(tm1108)* and *eat-3(ad426)* compared to *spg-7(ad2249)*, we conclude that defects in mitochondrial dynamics, to some extent, are suppressed or enhanced by the same pathways. Moreover, the suppressors of *fzo-1(tm1133)*-induced UPR^mt^ that were sorted into the functional groups ‘ribosome biogenesis’, ‘RNA processing’ and ‘translation’, reproduced comparably well in all secondary screens. Thus, attenuation of cytosolic translation may either be a general mechanism to suppress UPR^mt^ or, as discussed above, interfere with reporter expression. Among the enhancers, genes that sorted into the functional groups ‘ETC assembly factors’, ‘mitochondrial ribosome biogenesis’ and ‘mitochondrial translation’ showed the highest overlap among the secondary screens (Table S1), which demonstrates that disruption of mitochondrial translation robustly enhances UPR^mt^, independent of genetic background.

Twelve candidates that suppressed UPR^mt^ in the primary screen using *fzo-1(tm1133)*, enhanced UPR^mt^ in one or more of the secondary screens. Conversely, ten enhancers of *fzo-1(tm1133)*-induced UPR^mt^ suppress UPR^mt^ in at least one of the mutants in the secondary screens (listed in ‘opposing UPR^mt^ phenotypes’ sheet in Table S1). For example, knock-down of *icd-1*^βNAC^ suppresses P*_hsp-6_* _mtHSP70_*gfp* in all mitochondrial dynamics-related backgrounds, but enhances *spg-7(ad22449)*-induced UPR^mt^. Knock-down of *icd-1*^βNAC^ in *C. elegans* has been reported to induce UPR^ER^ in wild-type embryos (Arsenovic *et al.*, 2012). Furthermore, *icd-1*^βNAC^ has been described as a cytosolic stress sensor, which in the absence of stress associates with ribosomes to promote cytosolic translation, and which upon heat stress acts as a chaperone in the cytosol (Kirstein-Miles *et al.*, 2013). We recently showed that *icd-1*^βNAC^ is also a negative regulator of autophagy and that increased autophagic flux fuels mitochondria with certain triacylglycerols, thereby suppressing UPR^mt^ in both *fzo-1(tm1133)* and *drp-1(tm1108)* mutants (Haeussler *et al.*, 2020). Thus, blocking mitochondrial dynamics may reduce the flux of lipids into mitochondria, which can be compensated by the induction of autophagy. (We speculate that this may also apply to *eat-3(ad426)* mutants.) Conversely, we speculate that defects in mitochondrial homeostasis induced by a point mutation in *spg-7*, may exert stress to the cytosol and that this is normally compensated by factors, such as *icd-1*^βNAC^. Knocking-down *icd-1*^βNAC^ may therefore increase cytosolic stress, which in turn enhances UPR^mt^ in *spg-7(ad2249)* mutants. Taking the candidates into account that have opposing UPR^mt^ phenotypes in the secondary screens, 96% of the suppressors and 62% of the enhancers reproduce in *drp-1(tm1108)*, while 79% of the suppressors and 55% of the enhancers reproduce in *eat-3(ad426)*. We found the lowest overlap of candidate genes in *spg-7(ad2249)* mutants, with 59% of the suppressors and 49% of the enhancers reproducing in this background. Taken together, the results of the secondary screens demonstrate that there are candidates that when depleted act to influence UPR^mt^ signaling in general, whereas other candidates are specific to a certain type of UPR^mt^ induction, such as the disruption of mitochondrial dynamics.

### Transcription factor enrichment analysis identifies factors with roles in development, metabolism and oxidative stress response

Next, we identified TF binding sites in the promoters of our candidates using ChIP-seq datasets from the modENCODE project (Celniker *et al.*, 2009) in order to test for enrichment of TFs that bind to these sites. To that end, we used g:Profiler, a tool for functional enrichment analysis using over-representation (Raudvere *et al.*, 2019), which utilizes TRANSFAC resources (Knüppel *et al.*, 1994; Matys *et al.*, 2006). Using this approach, we found 15 TFs to be enriched in a statistically significant manner. Ten of these TFs only bind promotor regions of suppressors (7) or enhancers (3) (‘suppressor- or enhancer specific’). The remaining five TFs bind to promotor regions of both suppressors and enhancers (‘shared’) (Figure 3 and Table S3). The ‘shared’ TFs have previously been implicated in cell fate determination or developmental timing (Figure 3 and Table S3). Five out of seven ‘suppressor specific’ TFs have been shown to exclusively control developmental processes. The remaining two ‘suppressor-specific’ TFs are ELT-3^GATA3,4^ and HLH-11^TFAP4^, which have been shown to play a role in development, ageing and the response to oxidative stress (Gilleard *et al.*, 1999; Budovskaya *et al.*, 2008; Hu *et al.*, 2017) or to act as a dietary sensor that regulates metabolic gene expression, respectively (Soo-Ung *et al.*, 2009; Watson *et al.*, 2013).

**Figure 3:**
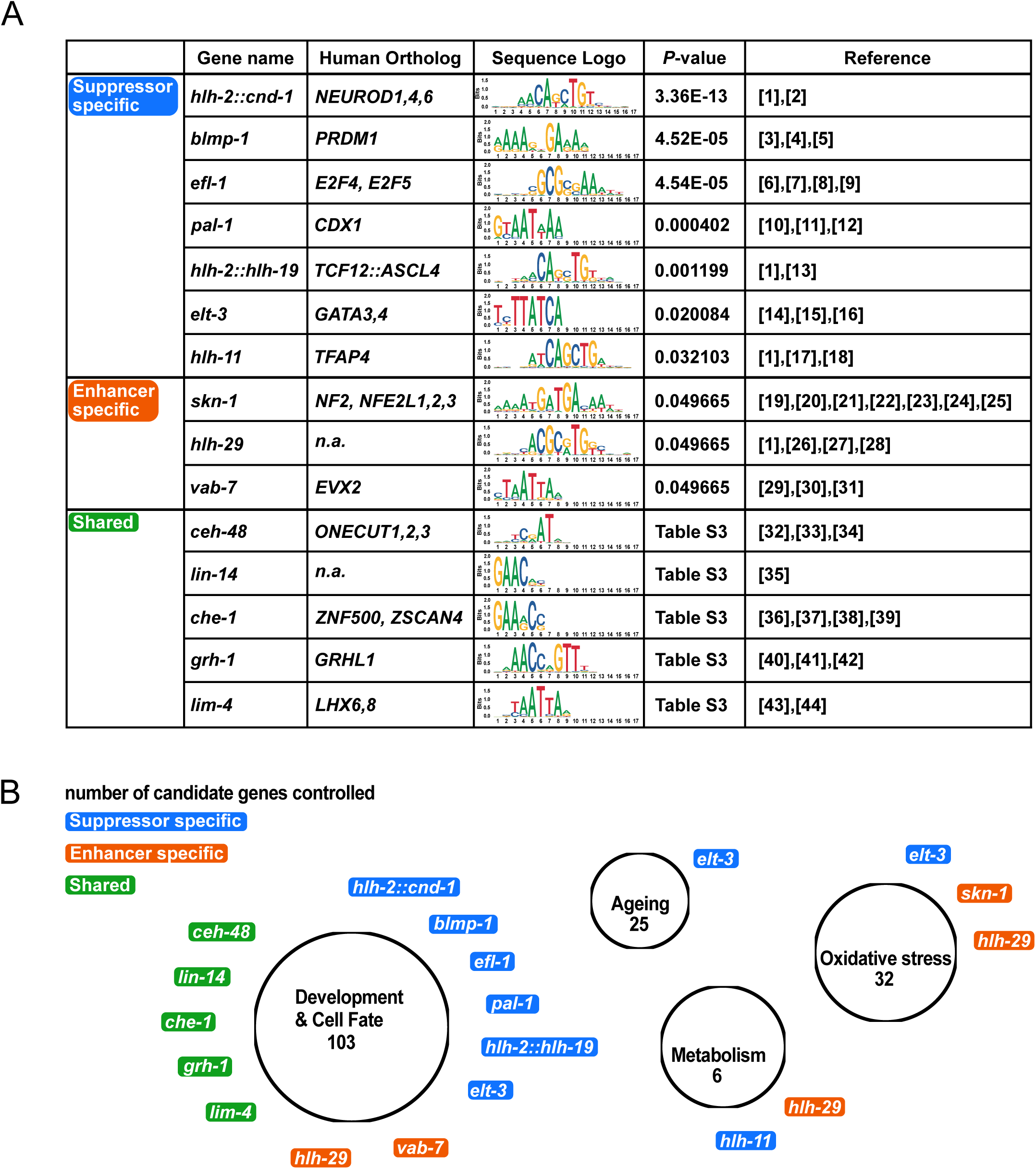
Enrichment analysis of transcription factors binding to promotors of candidate genes that suppress or enhance *fzo-1(tm1133)*-induced UPR^mt^. **(A)** Transcription factor (TF) binding sites were identified using the modENCODE database (Celniker *et al.*, 2009) and enrichment analysis was performed separately for suppressors and enhancers of *fzo-1(tm1133)*-induced UPR^mt^ using g:profiler (Knüppel *et al.*, 1994; Raudvere *et al.*, 2019). TFs that are statistically enriched among the candidate genes are shown. References: [1] (Grove *et al.*, 2009), [2] (Hallam *et al.*, 2000), [3] (Horn *et al.*, 2014), [4] (Huang *et al.*, 2014), [5] (Armakola and Ruvkun, 2019), [6] (Ceol and Horvitz, 2001), [7] (Garbe *et al.*, 2004), [8] (Chi and Reinke, 2006), [9] (Miller *et al.*, 2016), [10] (Baugh *et al.*, 2005), [11] (Maduro *et al.*, 2005), [12] (Lei *et al.*, 2009), [13] (Schwarz *et al.*, 2012), [14] (Gilleard *et al.*, 1999), [15] (Budovskaya *et al.*, 2008), [16] (Hu *et al.*, 2017), [17] (Soo-Ung *et al.*, 2009), [18] (Watson *et al.*, 2013), [19] (An and Blackwell, 2003), [20] (An *et al.*, 2005), [21] (Inoue *et al.*, 2005), [22] (Nargund *et al.*, 2012), [23] (Nargund *et al.*, 2015), [24] (Kim and Sieburth, 2018), [25] (Wu *et al.*, 2018) [26] (Neves and Priess, 2005), [27] (McMiller *et al.*, 2007), [28] (Quach *et al.*, 2013), [29] (Ahringer, 1996), [30] (Esmaeili *et al.*, 2002), [31] (Pocock *et al.*, 2004), [32] (Jacquemin *et al.*, 2003), [33] (Furuno *et al.*, 2008), [34] (Klimova *et al.*, 2015), [35] (Ambros and Horvitz, 1984), [36] (Chang *et al.*, 2003), [37] (Uchida *et al.*, 2003), [38] (Etchberger *et al.*, 2007), [39] (Rahe and Hobert, 2019), [40] (Huang *et al.*, 1995), [41] (Wilanowski *et al.*, 2002), [42] (Venkatesan *et al.*, 2003), [43] (Pradel *et al.*, 2007), [44] (Kim *et al.*, 2015). **(B)** Graphical representation of enriched TFs and the cellular processes they control. ‘Suppressor specific’ TFs are indicated in blue, ‘enhancer specific’ TFs in orange and ‘shared’ TFs in green. The number of candidate genes controlled by a certain group of TFs is indicated in each circle below the functional group name.

Three TFs (SKN-1^NFE2,NFE2L1,2,3^, HLH-29 and VAB-7^EVX2^) were identified to be ‘enhancer-specific’ (Figure 3 and Table S3). VAB-7^EVX2^ and HLH-29 are both required for certain aspects of development (Ahringer, 1996; Esmaeili *et al.*, 2002; Pocock *et al.*, 2004; Neves and Priess, 2005; McMiller *et al.*, 2007; Grove *et al.*, 2009) and HLH-29 has additional roles in fatty acid metabolism and energy homeostasis (McMiller *et al.*, 2007; Quach *et al.*, 2013). Furthermore, HLH-29 and SKN-1^NFE2,NFE2L1,2,3^ are regulators of the oxidative stress response (An and Blackwell, 2003; An *et al.*, 2005; Inoue *et al.*, 2005; Quach *et al.*, 2013) and SKN-1^NFE2,NFE2L1,2,3^ has previously been implicated in the UPR^mt^ pathway in *C. elegans* (Nargund *et al.*, 2012; Nargund *et al.*, 2015; Wu *et al.*, 2018).

In summary, we identified several TFs that have previously been implicated in oxidative stress response, cellular metabolism and development in *C. elegans*. Interestingly, *fzo-1(tm1133)* mutants have previously been shown to be slightly sensitive to oxidative stress and have increased levels of carbonylated proteins, a measure for oxidative damage (Yasuda *et al.*, 2011). Moreover, in *isp-1(qm150)* and *clk-1(qm30)* mutants, both of which have increased levels of reactive oxygen species (ROS) (Van Raamsdonk *et al.*, 2010; Yang and Hekimi, 2010; Dues *et al.*, 2017), UPR^mt^ activation has been shown to lead to ATFS-1^ATF4,5^-dependent expression of genes required for detoxification of reactive oxygen species (Wu *et al.*, 2018). This induction is orchestrated by ATFS-1^ATF4,5^ but may, to some extent, additionally be facilitated through activation of ELT-3^GATA3,4^ and HLH-29, as it has previously been shown for SKN-1^NFE2,NFE2L1,2,3^ (Nargund *et al.*, 2012; Nargund *et al.*, 2015; Wu *et al.*, 2018). The identification of many TFs controlling developmental processes is in agreement with our finding that GO-terms related to developmental processes are enriched among our dataset. This again highlights that the activity levels of critical cellular processes and responses in somatic tissues appear to be set during development. Finally, we previously found that the induction of autophagy suppresses UPR^mt^ in *fzo-1(tm1133)* mutants most likely through increased metabolic activity (Haeussler *et al.*, 2020). In our analysis, we identified two TFs, which regulate energy homeostasis and metabolic gene expression. This supports the notion that UPR^mt^ in *fzo-1(tm1133)* mutants acts to compensate for metabolic defects. In summary, we identified several TFs with roles in development, oxidative stress response and metabolism that had previously not been implicated in UPR^mt^ signaling. These TFs may be specific to UPR^mt^ in *fzo-1(tm1133)* or have general roles in UPR^mt^ signaling.

### Interactome of UPR^mt^ reveals potential new regulators

In order to determine whether any of the suppressors or enhancers that we identified have previously been shown to interact with *fzo-1*^MFN1,2^ or its mammalian orthologs *MFN1* or *MFN2*, we built a gene network containing all known interactions of *fzo-1*^MFN1,2^ and its mammalian orthologs *MFN1* and *MFN2*. Using the interaction databases ‘string-db.org’, ‘IntAct’, ‘BioGRID3.5’, ‘Genemania’, ‘CCSB’ and ‘mentha’ (Warde-Farley *et al.*, 2010; Calderone *et al.*, 2013; Orchard *et al.*, 2014; Rolland *et al.*, 2014; Oughtred *et al.*, 2018; Szklarczyk *et al.*, 2018), we included genetic and physical interactions (but not predicted interactions or co-expression data) and uploaded them to the cytoscape software (Shannon *et al.*, 2003) to calculate a complete interaction network. The resulting network contains 38 genes and 67 interactions (Figure S2). None of the 10 interactors of *fzo-1*^MFN1,2^ in *C. elegans* was identified in our screen (turquois dots in Figure S2). Next, we manually annotated the *C. elegans* orthologs of 24 interactors of *Mfn1* or *Mfn2* in mammals (except FAF2, MAVS, TCHP, SLC25A38 for which we did not find any orthologs in *C. elegans*, indicated in dark blue in Figure S2) but again did not find any overlap between the gene network and our screen dataset (orange dots in Figure S2). In summary, in our screen for modifiers of *fzo-1(tm1133)*-induced UPR^mt^, we did not find any previously known interactors of *fzo-1*^MFN1,2^. These could either have been missed in the RNAi screen, be essential in the *fzo-1(tm1133)* background or not have a function in mitochondrial homeostasis and, hence, UPR^mt^ signaling.

Similar to the approach described above, we used the 16 *C. elegans* genes currently associated with the GO-term ‘mitochondrial unfolded protein response’ (GO:0034514) (referred to as ‘input genes’), identified their human orthologs and included known physical and genetic interactors from the interaction databases ‘BioGRID3.5’, ‘IntAct’ and ‘mentha’ (Calderone *et al.*, 2013; Orchard *et al.*, 2014; Oughtred *et al.*, 2018) to calculate an interaction network containing 2603 genes and 4655 interactions (Figures S3, Figure S4, Figure S5). In this ‘UPR^mt^ome’, we identified 129 genes (including the 16 ‘input genes’), 36 of which are enhancers and 77 of which are suppressors of *fzo-1(tm1133)*-induced UPR^mt^, with a total of 213 interactions (Figure 4 and Table S4). For the ‘input gene’ *atfs-1*^ATF4,5^, we found five interactors (*gtf-2F2*^GTF2F2^, *lin-54*^LIN54^, *rps-6*^RPS6^, *spr-2*^SET^, *tbp-1*^TBP^) that suppress *fzo-1(tm1133)*-induced UPR^mt^ and the gene products of four of these localize to the nucleus (Sopta *et al.*, 1989; Lichtsteiner and Tjian, 1993; Wen *et al.*, 2000; Thomas *et al.*, 2003; Harrison *et al.*, 2006; Tabuchi *et al.*, 2011). These could potentially facilitate or directly be involved in the transcription of UPR^mt^ effectors upon activation of the UPR^mt^ response. Moreover, for the ‘input gene’ *ubl-5*^UBL5^, we found four interactors that overlap with our dataset of suppressors, three of which are splicing factors (*pqbp-1.2*^PQBP1^*, sfa-1*^SF1^*, snr-3*^SNRPD1^) (Thomas *et al.*, 1988; Krämer, 1992; Arning *et al.*, 1996; Imafuku *et al.*, 1998; Kambach *et al.*, 1999; Mazroui *et al.*, 1999; Waragai *et al.*, 1999). Of note, HUB1, the ortholog of UBL-5^UBL5^ in *Saccharomyces pombe*, has been shown to interact with components of the spliceosome. Furthermore, the loss of *HUB1* results in reduced splicing efficiency of a variety of mRNAs (Wilkinson *et al.*, 2004). Thus, the identification of the splicing factor genes *pqbp-1.2*^PQBP1^*, sfa-1*^SF1^ and *snr-3*^SNRPD1^ in our dataset presents an interesting potential link between UPR^mt^ activation and pre-mRNA splicing via UBL-5^UBL5^. In addition, we identified *taf-4*^TAF4^, which encodes an associated factor of transcription factor TFIID, to interact with the ‘input gene’ *sphk-1*^SPHK1,2^ and to suppress *fzo-1(tm1133)*-induced UPR^mt^ upon knock-down. *taf-4*^TAF4^ has previously been shown to be required for life span extension in *isp-1(qm150)*, *clk-1(qm30)* and *tpk-1(qm162)* mutants (Walker *et al.*, 2001; Walker *et al.*, 2004; Khan *et al.*, 2013). Finally, we identified many genes interacting with the ‘input gene’ *bar-1*^JUP,CTNNB1^, which has previously been shown to be involved in cell non-autonomous propagation of UPR^mt^ signaling (Zhang *et al.*, 2018). Among these interactors is phospholipase C (*plc-1*^PLCE^), which enhances *fzo-1(tm1133)*-induced UPR^mt^ and plays a central role in the inositol triphosphate (IP_3_) signaling pathway (Clandinin *et al.*, 1998; Kariya *et al.*, 2004). In summary, we identified several genes in our dataset using gene network analysis that have previously not been known to play a role in UPR^mt^ signaling in *C. elegans*. The genes with roles in pre-mRNA splicing and IP_3_ signaling may be particularly interesting in this respect. Furthermore, we propose that these genes may directly influence UPR^mt^ signaling through interactions with known players of the UPR^mt^ pathway.

**Figure 4:**
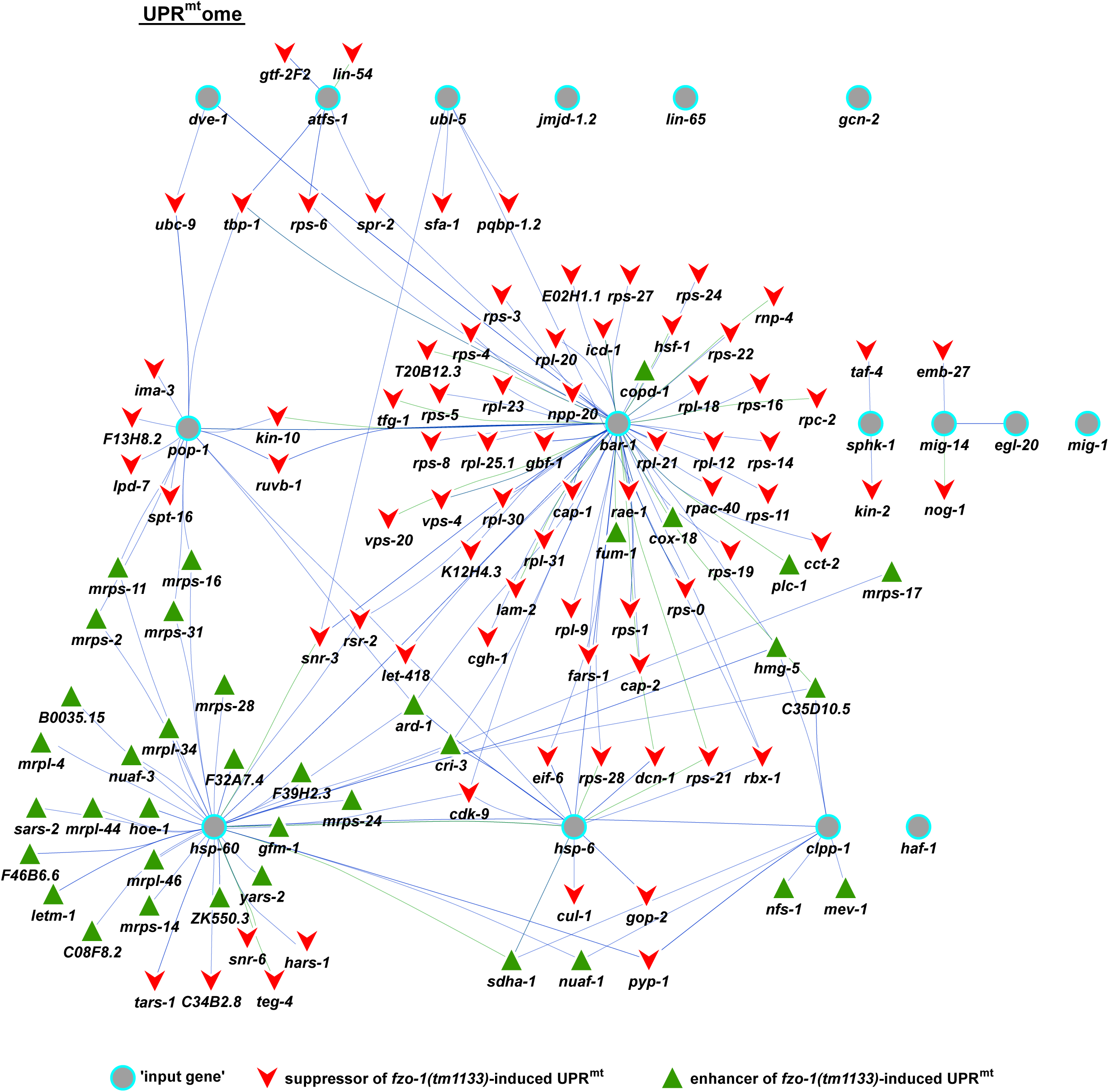
Analysis of a gene network – the UPR^mt^ome. Interactors of all genes that are currently associated with the GO-term ‘mitochondrial unfolded protein response’ and of their human orthologs were identified to build the complete UPR^mt^ome using ‘IntAct’, ‘BioGRID3.5’ and ‘mentha’ databases (Calderone *et al.*, 2013; Orchard *et al.*, 2014; Oughtred *et al.*, 2018). 129 genes are depicted, which overlapped between the complete UPR^mt^ome and the candidate list of our screen in *fzo-1(tm1133)* mutants. Turquois circles: ‘input genes’ currently associated with GO-term ‘mitochondrial unfolded protein response’, red arrowheads: suppressors of *fzo-1(tm1133)*-induced UPR^mt^ that overlap with the complete UPR^mt^ome, green triangles: enhancers of *fzo-1(tm1133)*-induced UPR^mt^ that overlap with the complete UPR^mt^ome. Interactions of two genes that were identified for *C. elegans* genes are indicated with green lines, interactions that were identified in human orthologs are indicated with blue lines.

### Interactome analysis reveals involvement of IP_3_ signaling pathway in UPR^mt^ regulation in *fzo-1(tm1133)*

In our gene network analysis, we identified *plc-1*^PLCE^, which encodes phospholipase C, as an interactor of *bar-1*^β-catenin^ (Byrne *et al.*, 2007). Interestingly, we and others found several genes that play a role in inositol triphosphate (IP_3_) signaling (Figure 5) (Liu *et al.*, 2014). The IP_3_ pathway is well known for its role in the regulation of intracellular calcium levels and transmits signals from the extracellular space via GPCRs and second messengers to the ER (Berridge, 2009). Thus, this signaling pathway may have a role in cell non-autonomous propagation of UPR^mt^. We identified the enzyme CDGS-1^CDS1^, which is essential for the production of phosphatidylinositol (PI) (Wu *et al.*, 1995; Vance, 1998), and EFR-3^EFR3B^, which targets PI-4-kinase (PI4K) to the plasma membrane (Nakatsu *et al.*, 2012). Furthermore, we identified the sole type I PIP kinase in *C. elegans,* PPK-1^PIP5K1A^ (Weinkove *et al.*, 2008), which phosphorylates PI4P to form PI(4,5)P_2_ (Ishihara *et al.*, 1996; Loijens and Anderson, 1996). PLC-1^PLCE^ is activated via GPCR and hydrolyzes PI(4,5)P_2_ to generate the second messengers DAG and IP_3_, known regulators of several signal transduction pathways (Clandinin *et al.*, 1998; Kariya *et al.*, 2004). One mechanism that is dependent on IP_3_-signaling is the release of calcium from the ER (Clandinin *et al.*, 1998; Kariya *et al.*, 2004; Kovacevic *et al.*, 2013). Interestingly, the IP_3_ receptor at the ER, ITR-1^ITPR1^, has previously also been identified as a suppressor of antimycin-induced UPR^mt^ (Liu *et al.*, 2014). Thus, it is tempting to speculate that altering IP_3_ signaling influences cellular calcium signaling in *fzo-1(tm1133)*, thereby affecting mitochondrial homeostasis and consequently UPR^mt^ signaling. Moreover, we propose that the effect on UPR^mt^ signaling may be indirect since we previously showed that knock-down of mitochondrial genes controlling calcium homeostasis does not induce UPR^mt^ in wild type (Rolland *et al.*, 2019). Furthermore, we propose that *fzo-(tm1133)* mutants may be more sensitive to changes in IP_3_ signaling and, hence, calcium signaling since these mutants may have altered ER-mitochondria contact sites, as shown for tissue culture cells lacking the mammalian ortholog MFN2 (de Brito and Scorrano, 2008; Cosson *et al.*, 2012; Filadi *et al.*, 2015, 2016; Leal *et al.*, 2016; Naon *et al.*, 2016; Basso *et al.*, 2018).

**Figure 5:**
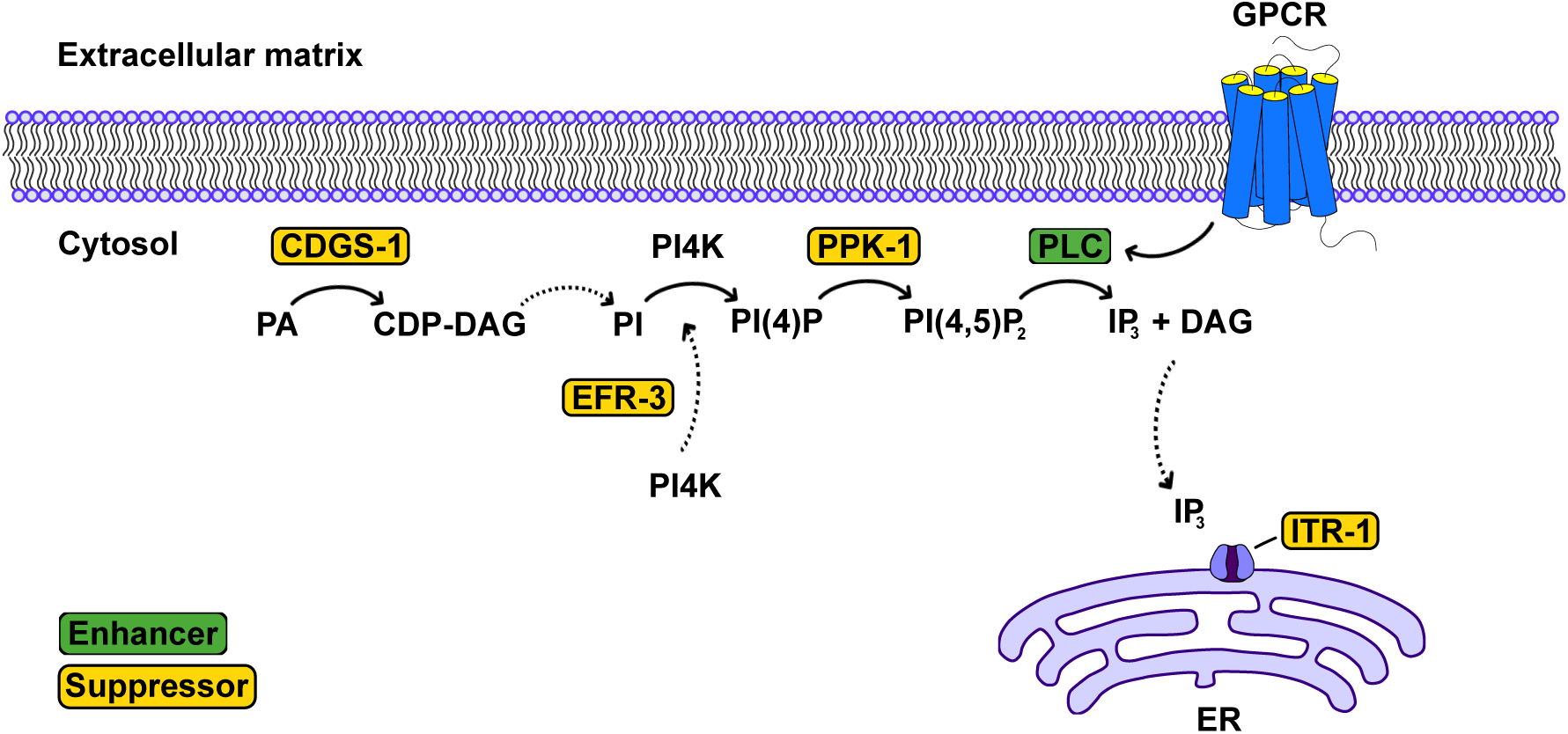
Candidate genes with roles in IP_3_ signaling. We identified four genes in our dataset that either play a direct role in the IP_3_ signaling pathway or are crucial for the synthesis of phosphatidylinositol-4,5-biphosphate (PI(4,5)P_2_). The IP_3_ receptor has previously been identified (Liu *et al.*, 2014). Suppressors are shown in yellow boxes, enhancers in green boxes. PA phosphatidic acid, CDP-DAG cytidine biphosphate-diacylglycerol, PI phosphatidylinositol, PI(4)P phosphatidylinositol-4-phosphate, IP_3_ inositol triphosphate, ER endoplasmic reticulum, GPCR G-protein coupled receptor.

### *miga-1(tm3621)* mutants show mitochondrial fragmentation and induce UPR^mt^

One of the enhancers we identified is *K01D12.6*, which is conserved from *C. elegans* to humans. The *D. melanogaster* ortholog of this gene has previously been identified in a screen for genes, which when knocked-down induce degeneration of photoreceptor cells. Furthermore, it was shown to be required for the maintenance of mitochondrial morphology and, hence, named ‘*Mitoguardin*’ (Zhang *et al.*, 2016). Moreover, the two orthologs of this gene in mammals (*MIGA1*, *MIGA2*) were found to regulate mitochondrial fusion and to be critical for mitochondrial function in human tissue culture cells and in mice (Liu *et al.*, 2016; Zhang *et al.*, 2016; Liu *et al.*, 2017). Therefore, we named *K01D12.6* ‘*mitoguardin homolog-1* (*miga-1*)’. We verified UPR^mt^ induction using the P*_hsp-60_* _HSPD1_*gfp (zcIs9)* reporter in the *miga-1(tm3621)* mutant background (Figure 6A). On average, the induction of P*_hsp-60_* _HSPD1_*gfp* is higher in *miga-1(tm3621)* animals compared to *fzo-1(tm1133)* animals. Moreover, we tested the effects of *miga-1(tm3621)* on steady-state mitochondrial morphology, which, in *C. elegans*, is carried out using a transgene that produces mitochondrial matrix-targeted GFP under the control of a body wall muscle cell-specific promoter (P*_myo-3_* _MYH_*gfp^mt^*) (Labrousse *et al.*, 1999; Ichishita *et al.*, 2008; Rolland *et al.*, 2013). Whereas wild-type worms show a tubular network of mitochondria, *miga-1(tm3621)* mutants have a ‘fragmented mitochondria’ phenotype, which is less severe than that caused by the loss of *fzo-1* (Figure 6B). In summary and in line with previous observations in other organisms, we observe significant changes in mitochondrial morphology in *miga-1(tm3621)* mutants, which are accompanied by the induction of UPR^mt^.

**Figure 6:**
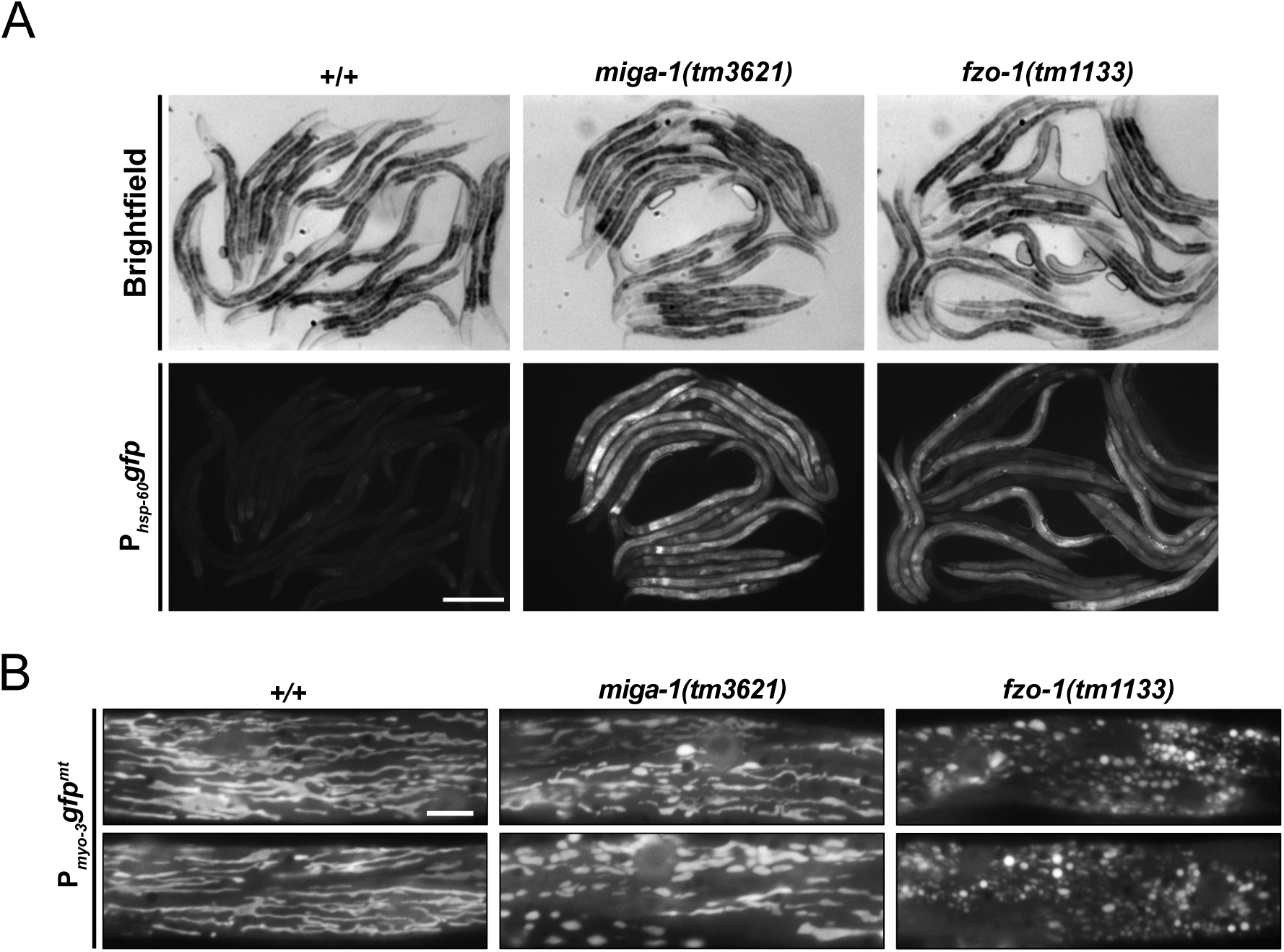
*miga-1(tm3621)* mutants induce UPR^mt^ and have altered mitochondrial morphology. **(A)** Fluorescence images of L4 larvae expressing P*_hsp-60_* _mtHSPD1_*gfp (zcIs9)* in wild type (+/+), *miga-1(tm3621)* or *fzo-1(tm1133)* mutants. Scale bar: 200 μm **(B)** Fluorescence images of L4 larvae expressing mitochondrial targeted *gfp* (P*_myo-3_gfp^mt^*) in wild type (+/+), *miga-1(tm3621)* or *fzo-1(tm1133)* mutants. Representative images are shown. Scale bar: 10 μm.

## ACKNOWLEDGEMENT

We thank members of the Conradt and Lambie labs and the ‘Mito Club’ for lively discussions. We thank M. Bauer, L. Jocham, N. Lebedeva and M. Schwarz for excellent technical support and S. Mitani (National BioResource Project, Tokyo, Japan) for *fzo-1*(*tm1133*), *drp-1(tm1108)* and *miga-1(tm3621)*. Some strains were provided by the CGC, which is funded by NIH Office of Research Infrastructure Programs (P40 OD010440). DFG funding (CO204/6-1 and CO204/9-1).

## SUPPLEMENTARY FIGURE LEGENDS

**Figure S1: Mutations in GTPases of the dynamin family induce the UPR^mt^.** *fzo-1(tm1133), drp-1(tm1108)* and *eat-3(ad426)* mutants differ in the level of induction of the P*_hsp-6_* _mtHSP70_*gfp (zcIs13)* reporter, as compared to wild type (+/+). *spg-7(ad2249)* is used as a positive control. Scale bar: 200 μm.

**Figure S2: Analysis of a gene network – the FZOome.** Interactors of *C. elegans* FZO-1 and of its human orthologs MFN1 and MFN2 were identified using ‘string-db.org’, ‘IntAct’, ‘BioGRID3.5’, ‘Genemania’, ‘CCSB’ and ‘mentha’ databases (Warde-Farley *et al.*, 2010; Calderone *et al.*, 2013; Orchard *et al.*, 2014; Rolland *et al.*, 2014; Oughtred *et al.*, 2018; Szklarczyk *et al.*, 2018). The identified candidates that suppressed or enhanced *fzo-1(tm1133)*-induced UPR^mt^ do not overlap with the FZOome. Turquois dots: interactors of FZO-1 in *C. elegans*; Orange dots: interactors of MFN1 or MFN2 which have orthologs in *C. elegans*; Blue dots: interactors of MFN1 or MFN2 in humans without any known orthologs in *C. elegans*.

**Figure S3: Subset of the UPR^mt^ome, direct interactors in *C. elegans*.** ‘Input genes’ associated with the GO-term ‘mitochondrial unfolded protein’ are indicated with turquois circles and their interactors in *C. elegans* are indicated with grey dots. Red arrowheads: suppressors of *fzo-1(tm1133)*-induced UPR^mt^ that overlap with the UPR^mt^ome subset; Green triangles: enhancers of *fzo-1(tm1133)*-induced UPR^mt^ that overlap with the UPR^mt^ome subset.

**Figure S4: Subset of the UPR^mt^ome, interactors of human orthologs.** ‘Input genes’ associated with the GO-term ‘mitochondrial unfolded protein’ are indicated with turquois circles and their interactors in humans are indicated with grey dots. Red arrowheads: suppressors of *fzo-1(tm1133)*-induced UPR^mt^ that overlap with the UPR^mt^ome subset; Green triangles: enhancers of *fzo-1(tm1133)*-induced UPR^mt^ that overlap with the UPR^mt^ome subset.

**Figure S5: Complete UPR^mt^ome.** ‘Input genes’ associated with the GO-term ‘mitochondrial unfolded protein’ are indicated with turquois circles and their interactors in *C. elegans* and humans are indicated with grey dots. Red arrows: suppressors of *fzo-1(tm1133)*-induced UPR^mt^ within the UPR^mt^ome; Green triangles: enhancers of *fzo-1(tm1133)*-induced UPR^mt^ within the UPR^mt^ome. Green lines indicate interactions in *C. elegans*, blue lines indicate interactions of human orthologs.

